# Engineering Threshold-Based Selection Systems

**DOI:** 10.1101/2021.02.04.429784

**Authors:** Katherine H. Pedone, Vanessa González-Pérez, Luciana E. Leopold, Channing J. Der, Adrienne D. Cox, Shawn Ahmed, David J. Reiner

**Affiliations:** Lineberger Comprehensive Cancer Center, University of North Carolina, Chapel Hill, NC 27599, USA; Curriculum in Genetics and Molecular Biology, University of North Carolina, Chapel Hill, NC 27599, USA; the Departments of Biology, University of North Carolina, Chapel Hill, NC 27599, USA; the Departments of Pharmacology, University of North Carolina, Chapel Hill, NC 27599, USA; the Departments of Radiation Oncology, University of North Carolina, Chapel Hill, NC 27599, USA; Institute of Biosciences and Technology, College of Medicine, Texas A&M Health Science Center, Houston, TX 77030, USA

**Keywords:** EGL-1, BH3-only, SMG-1, nonsense-mediated decay, NMD, 3’UTR, small GTPase

## Abstract

Using model organisms to identify novel therapeutic targets is frequently constrained by pre-existing genetic toolkits. To expedite positive selection for identification of novel downstream effectors, we engineered conditional expression of activated CED-10/Rac to disrupt *C. elegans* embryonic morphogenesis, titrated to 100% lethality. The strategy of engineering thresholds for positive selection using experimental animals was validated with pharmacological and genetic suppression and is generalizable to diverse molecular processes and experimental systems.

---

Harnessing model invertebrates to screen for small molecule inhibitors or for new genetic components of known processes is desirable because of the phylogenetic conservation of many key metazoan proteins and the infrequency of gene redundancy compared to mammals. Direct, unbiased screening for phenotypes is potentially powerful. Yet direct screening, whether with libraries of small molecules, RNAi, or chemical mutagenesis, can be difficult due to extensive phenotypic buffering and non-homolog redundancy in many biological processes. Additionally, potential screens depend on the availability of optimally suited reagents and/or phenotypes. These tools are frequently unavailable.

Screens can also demand sensitivity. For example, small molecule inhibitors must confer robust phenotypes, which may thus exclude potentially valuable lead compounds that would confer modest effects in initial screens. Screens can also be very laborious, particularly when scaled up to throughput levels necessary to detect rare positive candidates that confer incompletely penetrant or expressive phenotypes. Engineering of sensitized platforms that allow for identification of even atypical or rare candidates will help revolutionize approaches to contemporary experimental biology.

We have devised a method for engineering sensitized screening platforms using the experimental model organism *C. elegans*. This scheme is theoretically generalizable to any experimental organism. Our inspiration came from the activating mutation *let-60(n1046gf)* in the *C. elegans* ortholog of the human RAS oncoprotein. This G13E gain-of-function mutation induces ectopic 1^°^ vulval cells and hence a 100% penetrant Multivulva phenotype that is exquisitely sensitive to perturbation of downstream genes^1,2^, including genes that, when mutated alone, do not cause strong phenotypes, due to modulatory roles or redundancy^3-6^. Consequently, we aimed to develop a system where reagents similar to *let-60(n1046*gf) could be engineered on demand, and then exploited for genetic and pharmacological discovery screens.

We started with a signal known from mammalian studies and well conserved in *C. elegans*, but where the optimal genetic tools have not been previously generated by the research community. We used a conditional expression system whose full activation confers 100% toxicity. As needed, lethality could theoretically be conferred either at the level of failure of essential multicellular processes or the viability of single cells. This lethality thus establishes a threshold for positive selection for identifying suppressing compounds or mutations in the process of interest (**Fig. 1a**).

**Figure 1.**
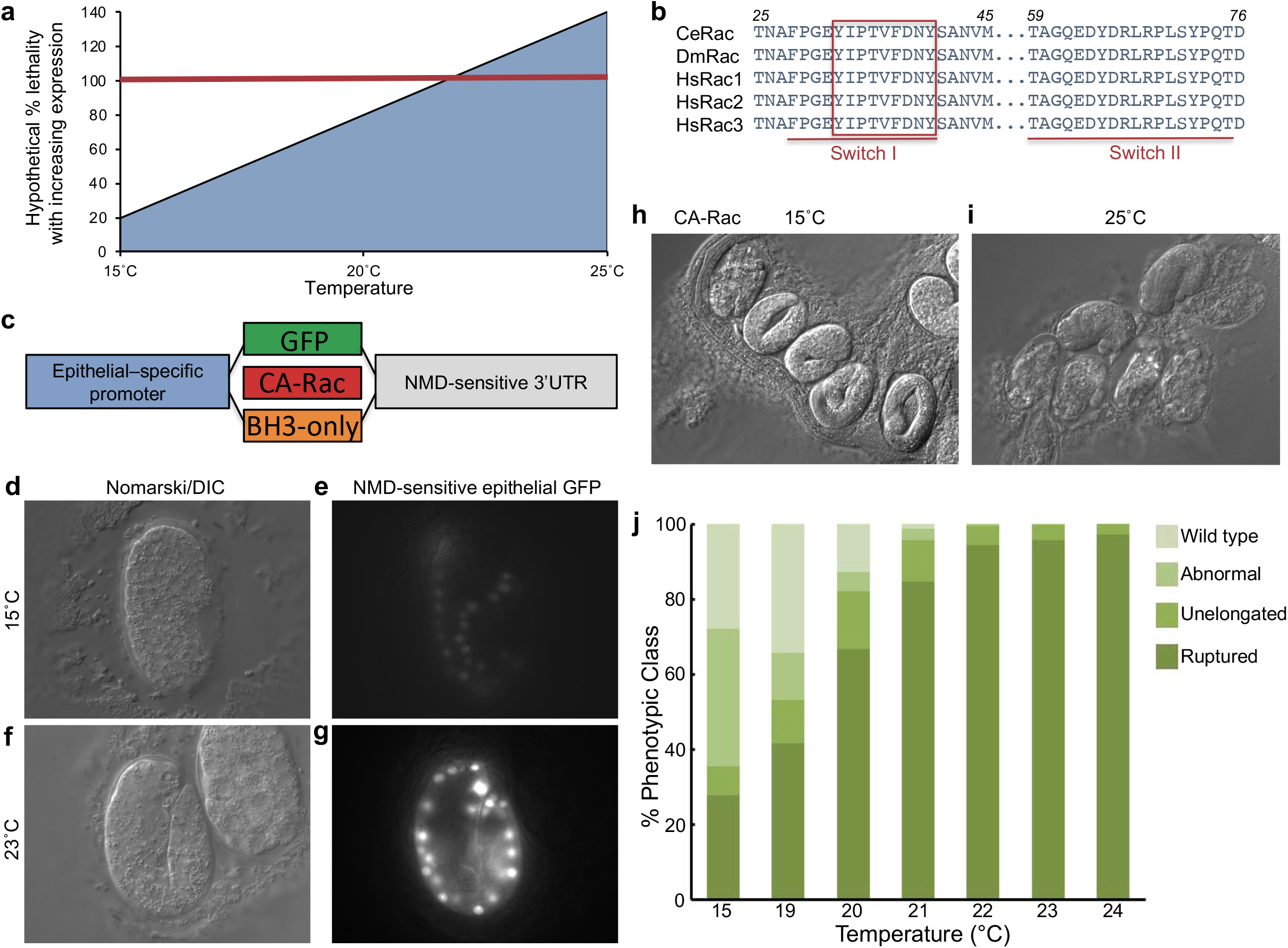
A system for conditional expression of signaling proteins to titrate to 100% functional threshold. a) A hypothetical graph of temperature-controlled levels of gene product required to reach the threshold of 100% lethality. **b)** 100% residue identity among Rac GTPases of *C. elegans, Drosophila melanogaster*, and humans in the structurally critical Switch I and II regions that harbor the core effector binding sequence (boxed). **c)** A schematic of plasmids for conditional expression of proteins, either control GFP, constitutively activated CED-10/Rac, or pro-apoptotic BH3-only protein EGL-1. The promoter is the “eFGHi” variant of the *lin-26* promoter, which drives expression in hypodermis (epithelial) cells in the embryo; the NMD-sensitive 3’UTR is inverted coding sequences from the *let-858* gene (A. Fire, pers. comm.). **d-g)** Temperature control of epithelial-specific expression from integrated transgene *reIs8* of GFP in epithelial cells under control of the hypodermal promoter and NMD-sensitive 3’UTR, in a *smg-1(cc546*ts) mutant background for temperature-sensitive perturbation of NMD. **d, e)** 100x DIC and epifluorescence micrographs, respectively, of a medial section of an enclosing embryo grown at 15^°^C reveals epithelial-specific expression and leakiness in the expression system. **f, g)** 100x DIC and epifluorescence micrographs, respectively, of medial sections of enclosed (center) and earlier stage (right) embryos grown at 23^°^C demonstrate elevated temperature-specific expression in epithelial cells and the absence of expression in earlier embryos, when *lin-26* expression is not activated and epithelial fate remains unspecified. See **Supplementary Figure 6** for hypodermal expression in different focal planes. **h**,**i)** 60x DIC images of *reIs6* animals expressing constitutively activated CA-Rac^CED-10^ at 15^°^C and 23^°^C, respectively. **h)** Animals grown at 15^°^C show a mixture of stages or hatched L1, and **i)** animals picked after growth for 24 hrs at 25^°^C show 100% rupture or arrested elongation. **j)** A curve of animal defects and survival at stepped temperatures from 15-24^°^C. Animals were binned into different classes based on morphology. “Abnormal” = observed lumps on the surface of hatched animals. “Unelongated” = intact embryos that failed to elongate. “Ruptured” = embryos that failed enclosure and so therefore exploded.

*C. elegans* Rac^CED-10^ is identical to human Rac in the critical Switch I and II regions involved in effector and regulator interactions (**Fig. 1b**). We focused specifically on essential morphogenetic functions of CED-10 that occur in differentiated cells that are post-mitotic, thus avoiding potentially complicated analyses of multiple biological processes in parallel^7^.

We expressed cDNAs behind the *eFGHi* variant of the *lin-26* promoter, which drives expression specifically in embryonic epithelial cells (**Fig. 1c; Fig. S1**)^8^. To achieve conditional expression, we used a synthetic 3’UTR that is subject to aggressive nonsense mediated mRNA decay (NMD), thus degrading the ectopically expressed mRNA. Transgenes were then expressed in a temperature-sensitive (ts) mutant for *smg-1*, an essential component of the NMD process.

We initially became interested in NMD by identification of a mutation in the gene *unc-97* that was suppressible by disruption of NMD via null (*re1* and *re861*) or temperature-sensitive (*cc545* and *cc546*) mutations in SMG-1, a conserved protein with a domain similar to that of PI3 Kinase. We confirmed the temperature-sensitivity of *smg-1* alleles *cc545* or *cc546* alleles using behavioral and semi-quantitative RT-PCR experiments for *unc-54(r293)*, which harbors a mutation in the 3’UTR of the endogenous *unc-54* gene that confers NMD-dependent loss of function and defective locomotion (**Figs. S2-5**). We found that *cc545* and *cc546* are predicted to cause single amino acid changes (**Fig. S6**), consistent with the hypothesis that temperature-sensitive mutations perturb protein structure or stability of SMG-1 in a manner that could allow for temperature-sensitive regulation of NMD.

We engineered expression of green fluorescent protein (GFP) cDNA under control of a synthetic NMD-sensitive (NMD^S^) 3’ UTR, all in the *smg-1(cc546*ts) genetic background. At restrictive temperature, where NMD is inactivated and hence mRNA stabilized and protein expressed, we observed high levels of GFP in embryonic epithelia. At permissive temperature, where NMD is functional, we observed lower levels of GFP, revealing some leakiness in NMD-dependent degradation of experimental mRNA (**Fig. 1d-g; Fig. S1**). Conditionally expressed GFP did not induce embryonic lethality, though weak morphological dysgenesis was observed (**Table S1; Fig. S7**). Thus, expression was temperature-sensitive, restricted to embryonic epithelial cells, and non-toxic.

To test our hypothesis that titratable expression of toxic proteins could be regulated with the NMD^S^ 3’UTR, we generated transgenic animals expressing Q61L mutant (constitutively active) CA-Rac^CED-10^. We observed that CA-Rac^CED-10^-dependent embryonic lethality was exquisitely titratable based on 1-degree steps of temperature. At restrictive temperatures we observed catastrophic failure of embryonic morphogenesis and elongation, with 100% embryonic lethality at 22^°^C. We decided to focus on 23^°^C as a fully penetrant non-permissive temperature for future experiments (**Fig. 1h-j**). Thus, our system is capable of generating conditional lethality calibrated to 100% lethality.

To further validate our expression system using a different signaling modality, we expressed the pro-apoptotic BH3-only protein, EGL-1^10^. At restrictive temperature, this transgene induced ∼100% lethality, was weakly suppressed by CED-3/caspase-directed RNAi, and strongly suppressed by the mutation *ced-3(n717)* in the C. elegans caspase (**Table S2**). At permissive temperature, little or no effect of this transgene was observed, thereby corroborating the threshold-based selection paradigm we sought. This temperature-sensitive NMD-sensitive system has also been harnessed for conditional expression of toxic signaling proteins for purposes of interrogating biological functions^11,12^.

We validated our screening methodology with mutations in Rac effectors and a selective small molecule inhibitor of mammalian Rac. The Pak serine/threonine kinase is a classic Rac effector that controls cytoskeletal dynamics and contributes to morphogenetic events downstream of Rac^CED-10^ in *C. elegans*^13,14^. Mutation of Pak^MAX-2^ partially suppressed CA-Rac^CED-10^-dependent lethality (**Fig. 2a**). Mutation of other known effectors was mostly inconsequential. Our genetic data indicate that CA-Rac^CED-10^ signals through at least one known effector.

**Figure 2.**
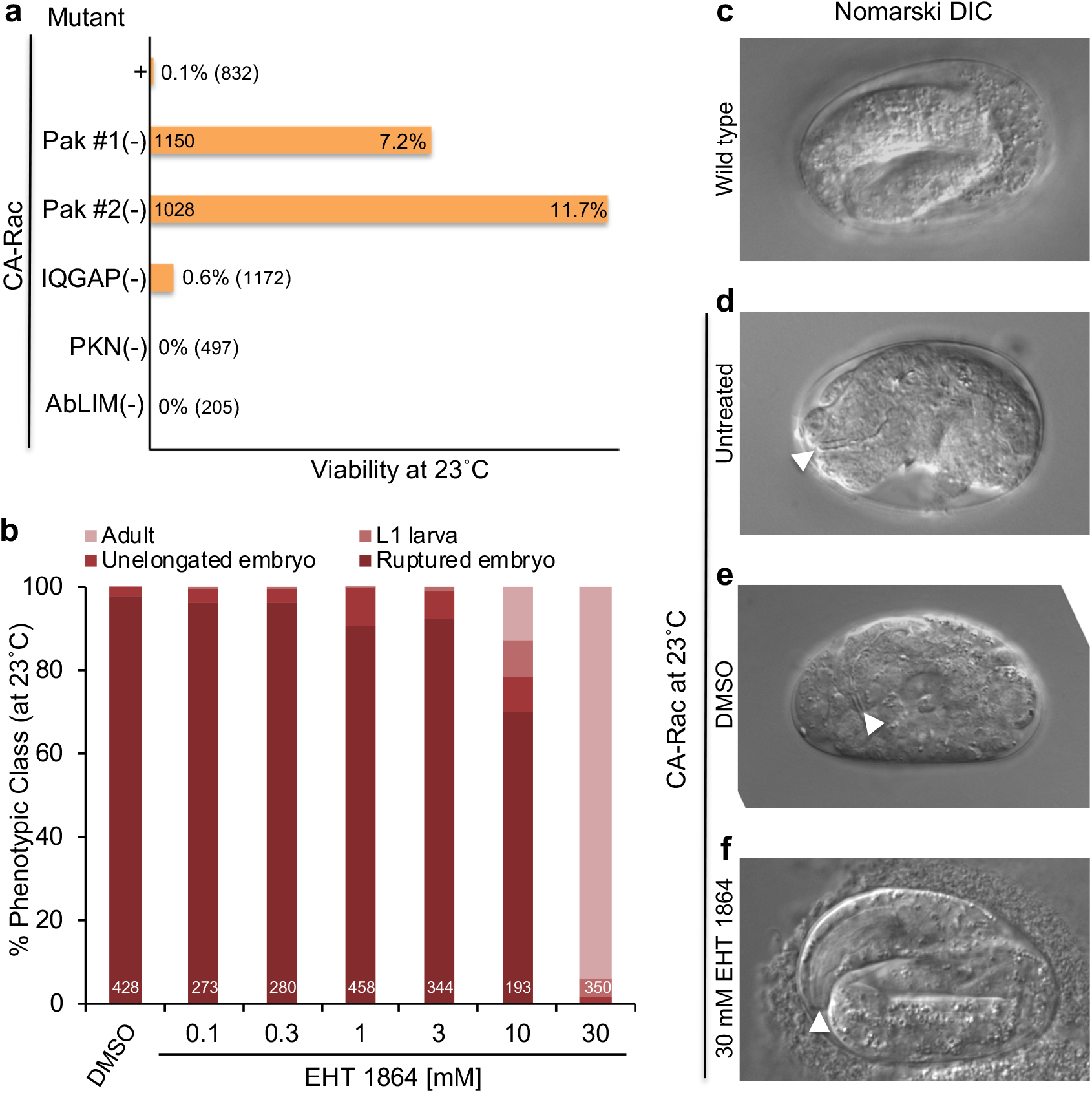
Genetic and pharmacological blockade of constitutively activated CED-10/Rac. a) In the *smg-1(cc546*ts); *s6[Plin-26::ced-10(Q61L)::NMD*^*S*^*3’UTR]* background, different mutations reduced levels of lethality. Two independent strain constructions with the *max-2(nv162)* mutation in the known Rac effector Pak suppressed lethality. Not shown is a synthetic lethal phenotype of *smg-1(cc546*ts); *reIs6* in combination with disruption of the other Pak ortholog, *pak-1(ok448)*. We constructed a strain with *pak-1(ok448)* as a heterozygote but could not homozygose the *pak-1* mutant chromosome. Unlike MAX-2/Pak, PAK-Pak has been implicated as an effector of both CED-10/Rac and CDC-42/Cdc42, as well as a GTPase- and kinase-independent component of the Pak-Pix-Git1 complex that regulates the cytoskeleton^13,14,17^. Thus, disruption of PAK-1, unlike disruption of MAX-is expected to impact multiple signaling systems, perhaps explaining the synthetic lethality observed when PAK-1 is deleted in the CA-CED-10/Rac transgenic background at any temperature. A putative null mutation in PES-7/IQGAP, *pes-7(gk123)*, a putative Rac effector identified in mammalian studies^18^, weakly suppressed lethality, while mutations in other Rac effectors PKN-PKN (*pkn-1(ok1673)*)^19^ and UNC-115/AbLIM (*unc-115(e2225)*)^20^ failed to suppress. **b)** Rescue of toxicity of constitutively active CED-10/Rac at higher concentrations of the Rac inhibitor, EHT 1864. Classes of phenotypes were binned as in **Figure 1. c)** DIC age of an embryo conditionally expressing GFP at 23^°^C. The lumen of the pharynx is out of the plane of focus, facing left. **d)** C image of a ruptured embryo conditionally expressing CA-Rac/CED-10 at 23^°^C. The lumen of the intact pharynx is in focus d facing left (white arrowhead), indicating that development persists even when epithelial morphogenesis is disrupted. **e)** DIC age of an embryo conditionally expressing constitutively activated Rac/CED-10 at 23^°^C and grown on 1% DMSO. The lumen of e intact pharynx is in focus and facing downward (white arrowhead). **f)** DIC image of an embryo conditionally expressing nstitutively activated Rac/CED-10 at 23^°^C and grown on 30 mM EHT 1864 in 1% DMSO.

We and others showed previously that the small molecule EHT 1864 blocked Rac effector signaling and induction of membrane ruffling in mammalian cells^15,16^. As overexpression of activated small GTPases like Rac could have unintended effects on cell biology, we asked if a selective inhibitor of Rac could suppress the effects of CA-Rac^CED-10^-dependent lethality. Treatment with EHT 1864 completely reversed the lethality conferred by CA-Rac^CED-10^ (**Fig. 2b-f; Fig. S8**). Most rescued animals appeared normal. Wild-type animals exposed to the same dose-response curve did not exhibit increased lethality. The structurally related negative control molecule EHT 8560 failed to rescue. These results affirm that it is CA-Rac^CED-10^ that conferred specific embryonic lethality and that compounds capable of inhibiting mammalian Rac can function to suppress Rac^CED-10^ activity in a distinct metazoan.

In summary, we have shown that it is possible to engineer specific biochemical pathways to establish the key threshold of 100% lethality. Such engineering can thereby selectively sensitize these pathways to both genetic and/or chemical suppression, via principles derived from classical suppressor genetics. We expect that the CRISPR revolution will further expand the flexibility, power and sensitivity of such engineered thresholds. We speculate that, if the appropriately sensitive tissue can be defined, this approach is generalizable to any experimental organism or biological process, including to humanized targets where this is advantageous. Conventional targeted therapies are based on the assumption that researchers have identified the best pharmacological target. A strength of our approach is that genetically sensitizing pathways in a model organism employs a different set of assumptions, and so lets the biology tell us which targets are most important for function. Thus, our study provides a paradigm for directed selection efforts targeting diverse pathways and various experimental genetic organisms in a manner that is broadly applicable in experimental biology.

## Acknowledgements

We thank M. Labouesse for pML433 and A. Fire for temperature-sensitive *smg-1* strains and plasmid pPD118.44, containing the synthetic NMD-sensitive (NMD^S^) 3’UTR, derived from the inverted *let-858* coding sequences. We thank Virginie Picard (ExonHit Therapeutics) for EHT 1864 and EHT 8560. This work was supported by NIH grant R01GM121625 to D.J.R. and R01CA175747 to C.J.D. K.H.P. was supported by NIH grant T32CA009156. Some strains were provided by the CGC, which is funded by NIH Office of Research Infrastructure Programs (P40 OD010440). Wormbase was used regularly.

## Supplementary information for

### Supplementary Methods

#### Strains and animal handling

Animals were cultured as described^1^ and handled in the default 20^°^C incubator if not otherwise noted, or in dedicated 15^°^C, 23^°^C or 25^°^C incubators. T-curves were performed in an incubator changed stepwise for each temperature. Strains used in this study are presented in **Supplementary Table 3**.

#### Microscopy

Animals were mounted in 2 mM tetramisole/M9 buffer on slides with agar pads. Animals were imaged with a Nikon Eclipse TE2000U microscope equipped with DIC optics, 40x, 60x and 100x oil objectives, with a DVC-1412 CCD camera (Digital Video Camera Company) controlled by Hamamatsu SimplePCI acquisition software. Some animal handling and imaging was performed on a Leica stereofluorescence microscope equipped with automated zoom optics.

#### Small molecule treatment

Pharmacological treatment with EHT 1864 and inactive analog EHT 8560 (provided by Virginie Picard, Exon Hit Therapeutics) was performed in 6-well microtiter plates with small volumes of agar, with only corner wells used. EHT 1864 was diluted to the appropriate concentration in 1% DMSO, with 1% DMSO without inhibitor as a control. Animals were grown in an incubator dedicated to 23^°^C, with all animals for a given assay grown in parallel.

#### Locomotion assay

Animals were subjected to a circumferential locomotion assay as described previously^2^. Briefly, young adult animals were picked to the center of a 10 cm plate seeded 1 day previously, the origin was marked on the bottom, and animals were allowed to roam freely for 20 min, at which point they were arrested by placing at -20^°^C for 5 min. The distance from the origin was measured for each animal.

#### Molecular biology

Details of plasmid construction and PCR detection of mutations is available upon request. Primers used in this study are presented in **Supplementary Table 4**. Plasmids used are presented in **Supplementary Table 5**.

### Supplementary Protocol

We provide the following protocol to generate reagents that impose conditional lethality. Many of the principles are universal, while some of the tools and reagents discussed here are specific for use in *C. elegans*. The details for conditional expression of toxic proteins are likely to vary in other systems.

#### Selection of biological processes to target

We targeted CED-10/Rac because of its high level of sequence and functional conservation. We expressed it specifically in epithelia undergoing morphogenesis because we wanted to avoid targeting cell fate decisions or cell proliferation. *The protein/pathway of interest to be targeted by each investigator is likely to be guided by their specific research interests. We emphasize that not all tissues or organisms may be optimally suited to develop this assay. Rather, we suggest that researchers develop assays using whichever system and/or tissue is best suited for the process they wish to target*.

- Proteins whose expression is to be controlled to confer toxicity should be selected based on potential for gain-of-function toxicity, either through constitutive activation through mutation (gain of normal function), mis-expression (gain of novel function), or removal of negative regulator through conditional knockout, including engineered temperature-sensitive mutations^6^ or chemogenetic tools like the auxin inducible degron^3-5^.
  - Mutational activation of CED-10/Rac via the Q61L mutation described here to disrupt morphogenesis is an example of gain of normal function.
  - Mis-expression of the EGL-1/BH3-only protein in epithelial (hypodermal) cells to induce apoptosis is an example of gain of novel function.
- Tissues to be targeted should be guided based on prior evidence of function in those tissues.
  - The case for the morphogenetic hypodermis as target tissue is that it is post-differentiation and post-mitotic^7^. CED-10/Rac, a well-known regulator of cytoskeletal dynamics, has also been validated to play an important role in a series of post-mitotic morphogenetic events in the *C. elegans* mid-embryo^8^.
  - Although cells in the embryonic hypodermis do not typically undergo apoptosis^9^, we chose to express EGL-1/BH3-only in this tissue so we could compare directly to toxicity conferred by mutationally activated CED-10/Rac, with only the cDNA expressed differing between the two tools. Indeed, we screened through far more candidate transgenes for EGL-1/BH3-only expression than for CED-10/Rac expression, and were never able to consistently obtain 100% lethality.
  - A more promising tissue in which to evoke apoptosis through ectopic expression of EGL-1/BH3-only would be neurons, among which a large number undergo apoptosis during the normal course of *C. elegans* development^9,10^.
- Piloting toxicity: Prior to assembling the entire system, a quick pilot experiment to evaluate the original premise may increase the likelihood of success. For CED-10/Rac, we overexpressed three small GTPases known from other systems to control cytoskeletal dynamics during cell movements – Rac, Rho and Cdc42^11^ – by generating high-copy transgenes with cosmid clones of genome intervals containing each gene. This is a relatively quick assay. Of these, CED-10/Rac conferred diverse defects in morphology with greater penetrance than did Rho/RHO-1 or Cdc42/CDC-42. This was our sole indicator that CED-10/Rac was a promising candidate.

#### Selection of conditional expression systems

*Like selection of proteins of interest, selection of tissues to be targeted should be performed in consultation with the literature and/or an expert in the field to make decisions most likely to result in success*.

- The promoter is part of the conditional expression system in our application, driving expression only in a defined tissue. In this context, we selected a specific variant of the *lin-26* promoter, eFGHi, previously demonstrated to express mainly in hypodermal cells during embryonic morphogenesis. The complete range of *lin-26* expression includes post-embryonic epithelial cells as well as diverse support cells, and thus would likely complicate interpretation and make subsequent analysis more difficult^12^. Other promoter types could target other tissues, developmental stages, or environmental conditions, including heat-shock promoters^13^.
- Both spatial and temporal control.
  - In addition to the eFGHi variant of the *lin-26* promoter to confer spatial specificity, we coupled conditional degradation through the 3’UTR to confer temporal specificity via temperature sensitive mutations that abrogate NMD. Nonsense-mediated decay has a robust history in *C. elegans*^14^ but has the potential downside of regulating many transcripts of the animal, including potentially those that are part of normal regulation of development of genome surveillance, rather than the aberrant premature termination codons for which NMD is best known^15^.
  - A similar informational suppressor system uses an intron from the *unc-52* gene that is selectively spliced by MEC-8, a splicing factor for which a temperature-sensitive allele exists. Retention of the intron abrogates gene function, and splicing of the intron relies on *mec-8(u218*ts)^16^.
  - We point to recent advances with the auxin-inducible degron (AID), a conditional degradation system that requires a substrate protein to be tagged with the AID sequence, requires the TIR1 co-factor that can be expressed in different tissues or different times, and requires addition of the auxin small molecule to trigger degradation^3^; Ashley *et al*., in press; preprint available at https://doi.org/10.1101/2020.05.12.090217.

### Generation of transgenes expressing toxic proteins

To engineer toxicity, one must be able to generate tools under conditions that permit viability. As noted in this study, the *smg-1(*ts) system is leaky: at 15^°^C, protein is expressed and causes some level of toxicity. This feature of the system made it difficult to generate conditionally toxic transgenes in the *smg-1(*ts) animals at 15^°^C: we systematically biased against the weakly expressing transgenes that could be tolerated by the animal. We therefore developed a protocol to reproducibly isolate toxic transgenes, starting with the wild-type animal in which expression was less leaky.

- Inject the DNA mix, including selection markers (P_*myo-2*_*::gfp* and/or *rol-6(*d)) into a wild-type background, and isolate scores of independently derived extrachromosomal arrays.
- Plate each candidate line on bacteria expressing dsRNA targeting a gene required for NMD (we used *smg-1*, clone C48B6.6, address I-3K02).
- Select transgenes causing a range of severity of defects when grown on bacteria expressing *smg-1*-directed dsRNA^17^. Score several lines semi-quantitatively (for speed) and forward for further analysis. (We do not freeze the scores of lines, typically settling for ∼5 for freezing and further analysis).
- We then integrate extrachromosomal arrays in the wild-type animal background using a variety of published protocols: UV, gamma irradiation, etc.^18^.
- Re-test with *smg-1(RNAi)*. Some integrated lines are approximately the same strength, while others are markedly weaker, often accompanied by decreased expression of the P_*myo-2*_*::gfp* pharyngeal GFP co-expression marker. We discard the latter.
- Outcross resulting integrants 4x into the wild-type strain background. Freeze the resulting outcrossed strain.
- For candidates of different levels of severity on *smg-1(RNAi)*, cross into the *smg-1(cc546*ts) strain, using either SNP-snip PCR detection of the *cc546* lesion (Supplementary Figure 6) or balancing the *smg-1* locus with a mutation in the closely linked *unc-87*. (*N*.*B*. most strains will display some lethality at permissive temperature of 15^°^C).
- Be extremely careful about genetic drift; we observed that strains conferring toxicity became less severe when cultured over many generations.
- To prevent drift, we perform the following steps.
  - Immediately starve and freeze candidate strains. We also test thaw and quantitatively assess lethality to ensure that the frozen strain has not drifted.
  - Maintain the strain as a starved, parafilmed plate for a few months. From this plate, we would extract animals weekly with a chunk of agar, to constantly provide “fresh,” undrifted animals for assays or further crosses.
  - Severity can be reset by reconstructing the strains by crossing in *unc-87* to balance *smg-1* and then re-isolating the original strain. Thus, we hypothesize that modifying mutations accumulate over time, though we cannot rule out silencing of the transgene.
- Assess toxicity for each *smg-1(*ts)+transgene combination for those reaching 100% lethality. Select for further analysis those that are not at the upper end of the temperature range; above 25^°^C, animals can be difficult to culture without drift.
- Generate an efficacy curve at a range of temperatures (T-curve). Procced only with those that confer 100% lethality at or below 25^°^ but are fecund at 15^°^C
- Validate source of toxicity by RNAi-dependent depletion of the toxic protein, small molecule inhibition, genetic perturbation of “downstream” intermediaries, etc. (see **Fig. 2**).

This protocol should yield strains with properties similar to those described here. However, we note that successfully attaining the desired goal of 100% lethality is a function of multiple variables. Selection of transgenes that confer the strongest defects biases the results towards success, but some systems may not be capable of reproducibly driving 100% lethality. As an example, we use our system with EGL-1/BH3-only expressed conditionally in hypodermal cells. The strain we analyzed was selected from many scores of candidates, and still fell short of reproducibly reaching 100% lethality. (We could attain 100% lethality at 27^°^C, but resultant animals were sickly and sterile).

## Supplementary Figure Legends

**Supplementary Figure 1.**
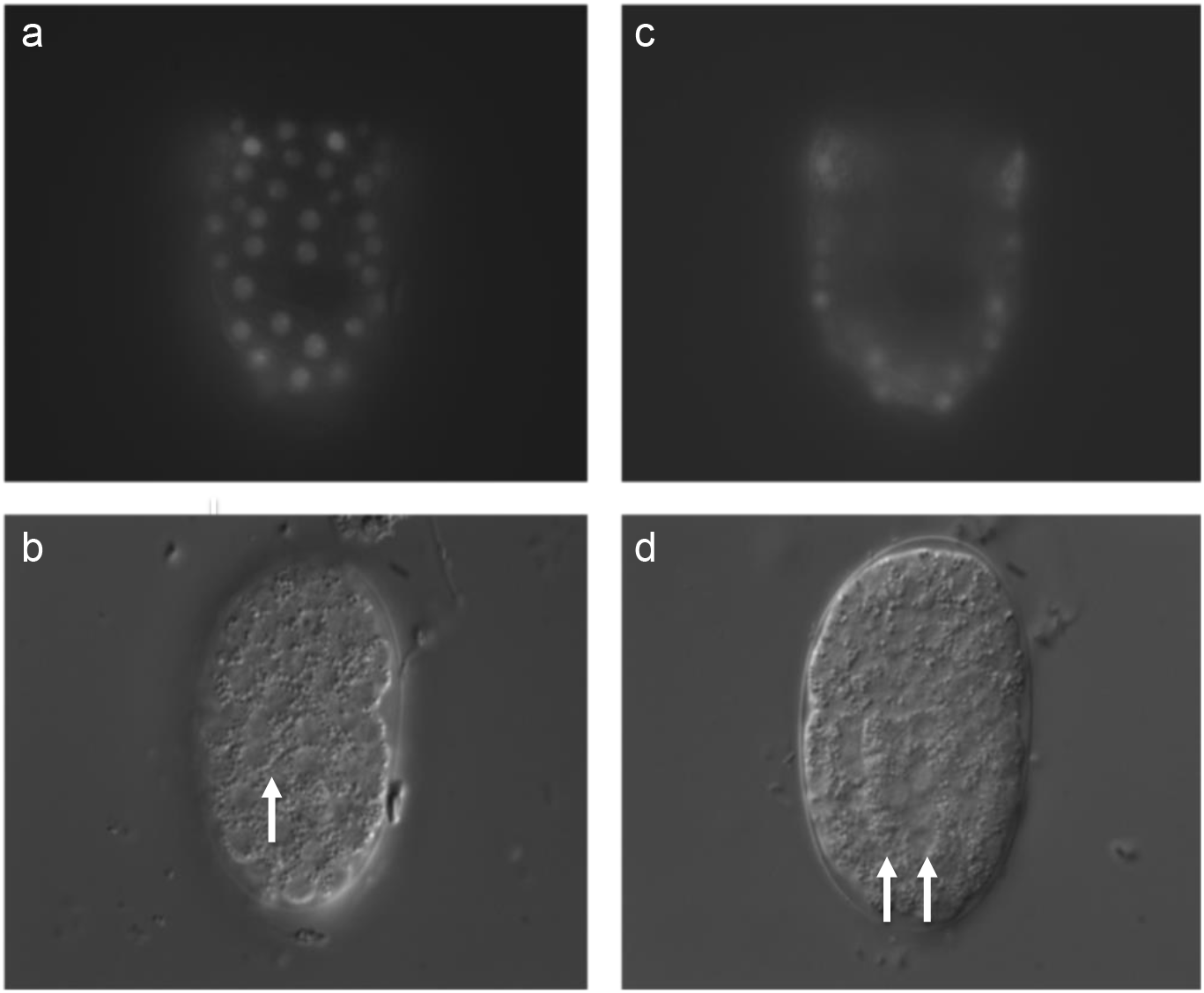
Epithelial-specific GFP expression at 15°C. a-d) The same early enclosure staged *smg-1(cc546*ts); *reIs8[P*_*lin-26*_*::gfp::NMD*^*S*^*3’UTR]* embryo in different focal planes. **a, b)** Epifluorescence (500 msec exposure) and DIC images, respectively, of the dorsal surface of the embryo, with arrows indicating a row of intercalating epithelial cells. **c**,**d)** Epifluorescence (500 msec exposure) and DIC images, respectively, of a medial section of the embryo, with arrows indicating the column of intestinal cells.

**Supplementary Figure 2.**
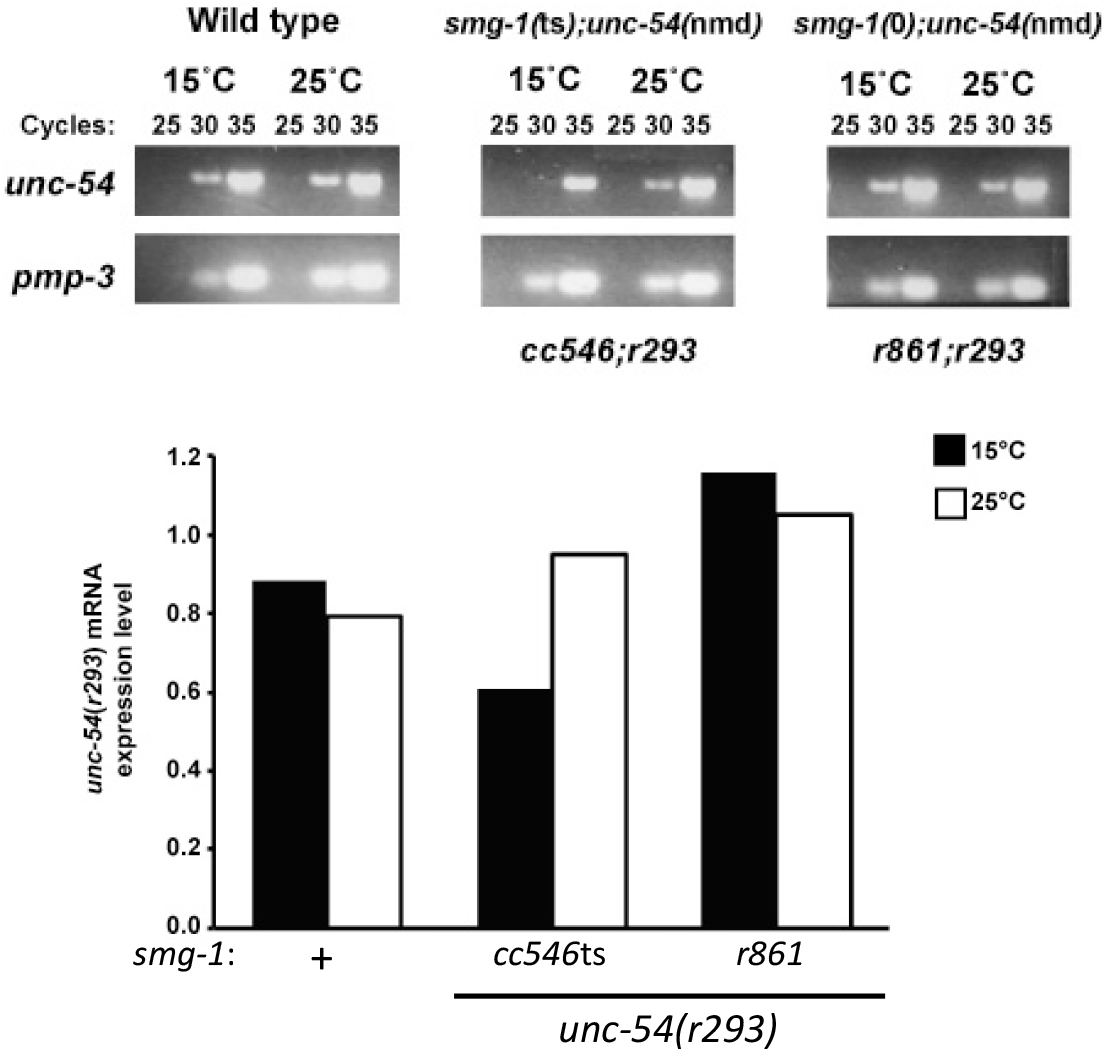
NMD-dependent differences in gene expression. Animals harboring the *cc546* temperature-sensitive mutation in *smg-1* have increased *unc-54(r293)* mRNA levels at 25^°^C by RT-PCR, with *pmp-3* RNA as a control. RNA extractions were performed on pools of adult animals raised at either 15^°^C or 25^°^C. cDNA preparations of each strain were subjected to 25, 30 or 35 cycles of PCR with *unc-54*-specific primers. Temperature-dependent differences were visible at 30 cycles with the *cc546*ts allele of *smg-1* used in this study but not the *smg-1(+)* or *smg-1(r861)* putative null mutation, as shown in the graph. Band intensities were quantified using the Image J gel analysis tool. Experiment was performed two times.

**Supplementary Figure 3:**
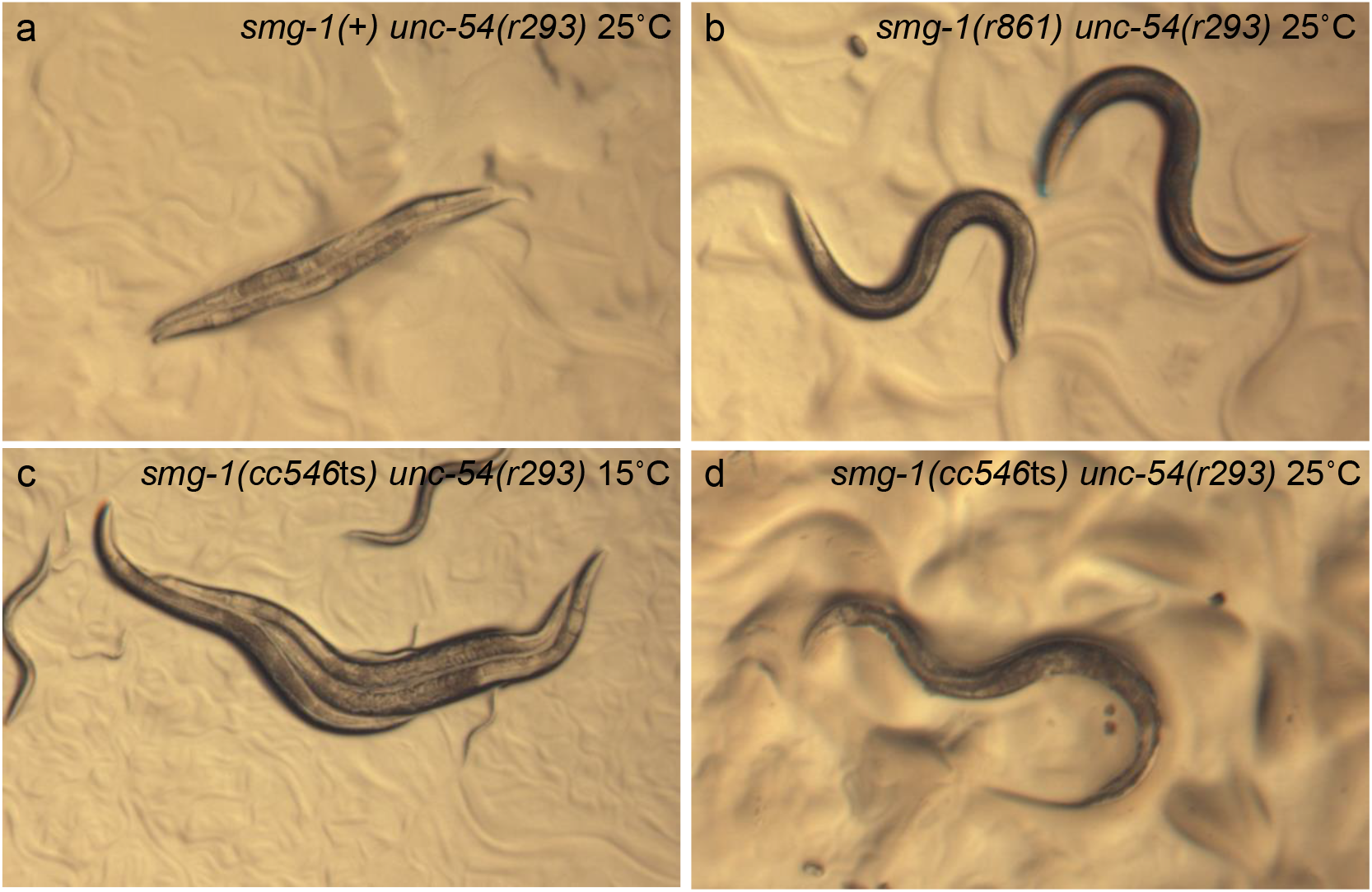
A *smg-1* temperature-sensitive allele regulates locomotion. All strain backgrounds harbor NMD-sensitive *unc-54(r293*). Photomicrographs were captured from agar plates with 25 msec exposures under same lamp settings. Body posture is representative of locomotion and hence myosin production by the *unc-54* gene and its NMD-sensitive mutation in the *unc-54* 3’UTR, *r293*: deep body bends represent typical locomotion, shallow bends represent flaccid paralysis. **a)** *unc-54(r293)* animals were paralyzed and egg-laying defective (Egl). **b)** The locomotion and Egl defects of the *r293* mutant were strongly rescued by loss of *smg-1* function. **c)** Locomotion and Egl defects were not as severe with *cc546*ts as with *smg-1(*+) at 15^°^C and **d)** are completely suppressed at 25^°^C, consistent with *cc546*ts being temperature sensitive.

**Supplementary Figure 4:**
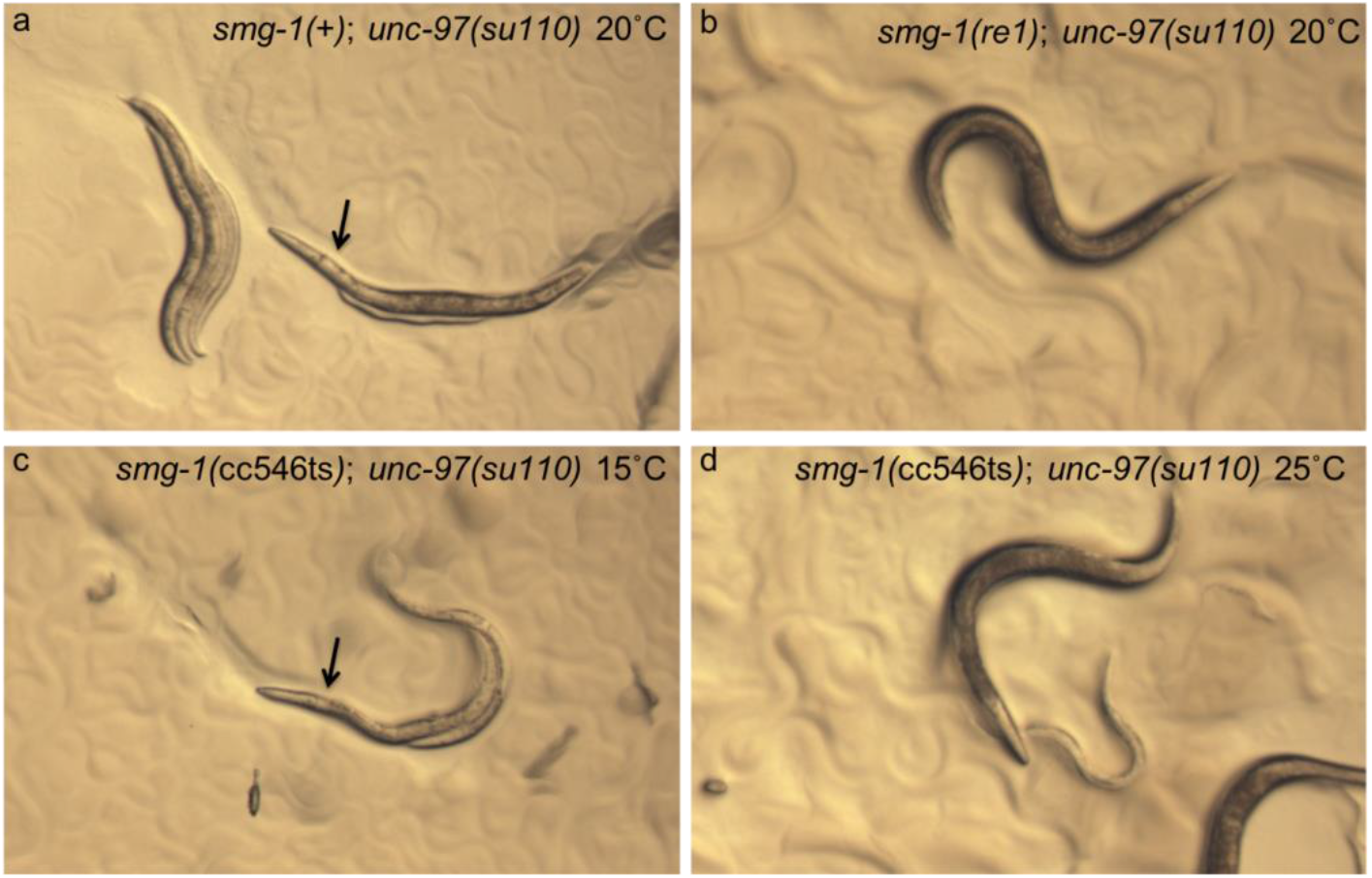
TS NMD-sensitive *unc-97(su110)*. Upon crossing into the reference strain for *unc-97(su110)*, HE110, we observed that the strain contained a background mutation partially suppressing the Unc phenotype of *unc-97(su110)*. Whole genome sequencing of this strain identified a nonsense mutation in *smg-1*, which we named *re1* (see **Supplementary Figure 6**).). Photomicrographs were captured from agar plates with 25 msec exposures under same lamp settings. Body posture is representative of locomotion and hence PINCH production by the *unc-97* gene and its NMD-sensitive mutation in the *unc-54* 3’UTR, *r293*: deep body bends represent typical locomotion, shallow bends represent flaccid paralysis. Arrows point to a clear area posterior to the pharynx that indicates a clear patch in the intestine that indicates distension with liquid due to defective defecation. *unc-97(su110)* animals alone are paralyzed, Egl, and constipated. **b)** These phenotypes are suppressed by the *smg-1(re1)* mutation crossed back into the *unc-97(su110)* background, **c)** not suppressed by *smg-1(cc546*ts) at 15^°^C but **d)** suppressed by *smg-1(cc546*ts) at 25^°^C. Mutants for *unc-97* have been reported to have mechanosensory defects (Chen and Chalfie, 2014), and are thereby sluggish and do not move on plate assays. Consequently, we did not include *unc-97(su110)* in our locomotion analysis for **Supplementary Figure 5**.

**Supplementary Figure 5.**
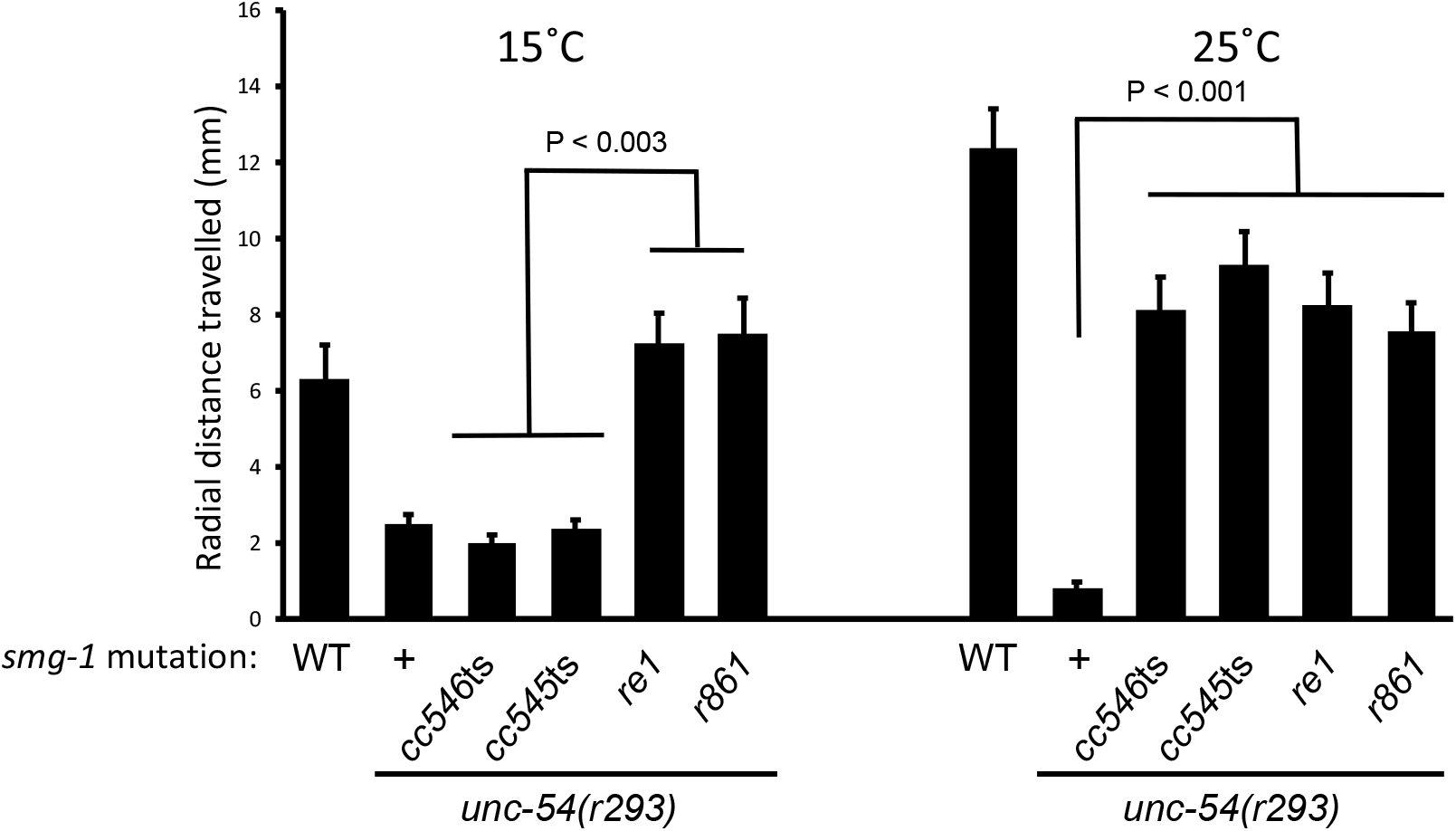
RT-PCR detection of NMD-dependent differences. Putative null mutations in *smg-1* suppress locomotion defects conferred by the aberrant *unc-54(re293)* 3’UTR at both 15^°^C and 25^°^C. *smg-1(cc545*ts) and *smg-1(cc546*ts) fail to rescue locomotion defects at 15^°^C but rescue at 25^°^C. All animals were scored in sequential assays on the same day, 20 minute assays, then transferred to -20^°^C for 5 minutes to arrest locomotion, then counted.

**Supplementary Figure 6:**
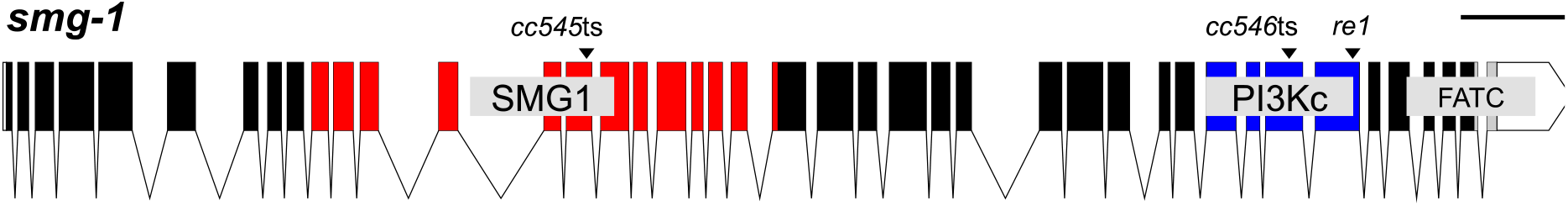
Identification of lesions in *smg-1*. The exon-intron boundaries of the *smg-1* gene are shown. Domains are SMG1 (red), PI3Kc (blue) and FATC (gray), UTRs are white. Scale bar = 1000 bp. *cc545ts* is an ACA>ATA transition in exon 15 that causes a T761I missense change. *cc546ts* is an unusual ATG>TTG transversion in exon 35 that causes a M1957L missense change. *re1* is an unusual GAG>TAG transversion in exon 36 that causes an E2093^*^ nonsense change. *cc546ts* is detectable by SNP-snip: restriction enzyme Msl I cuts the wild-type but not the mutant sequence.

**Supplementary Figure 7.**
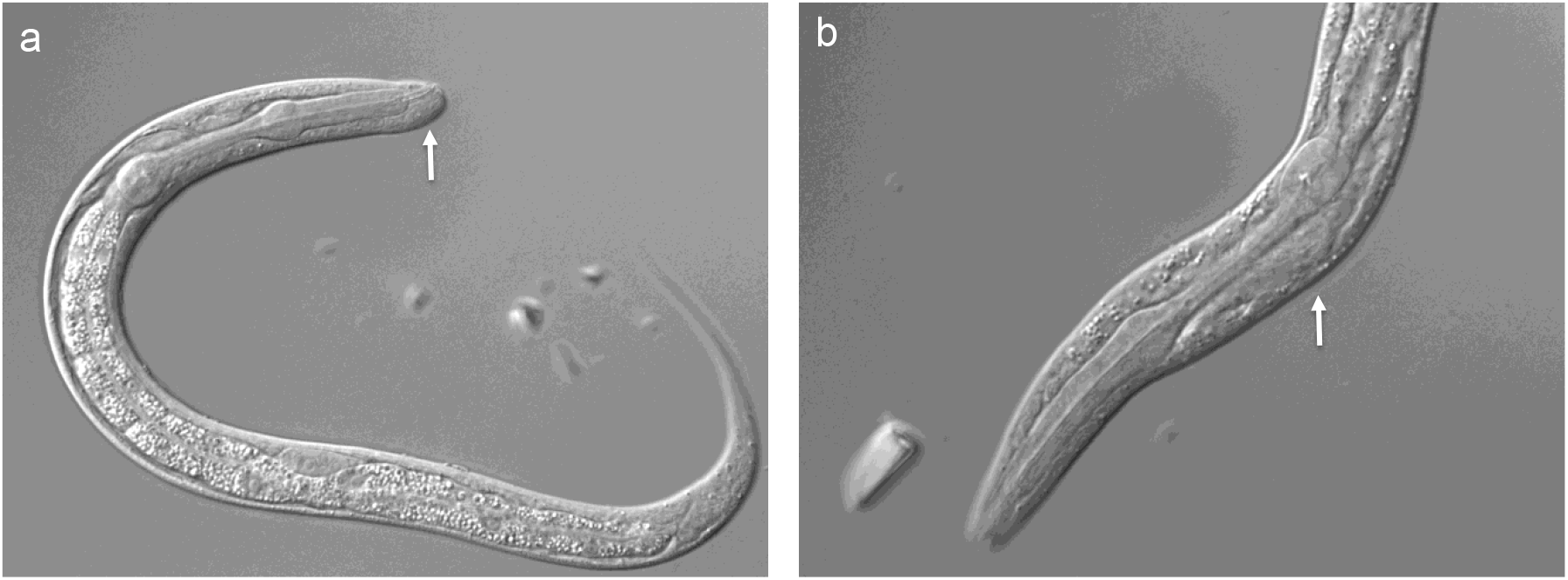
Weak morphogenetic phenotypes. Low penetrance and low expressivity phenotypes are caused by the GFP over-expressing strain *smg-1(cc546*ts); *reIs8[Plin-26::gfp::NMD*^*S*^*3’UTR]*, perhaps due to over-represented promoter sequences titrating factors important for morphogenesis. Arrows indicate mild bulges in the animal’s epithelium, frequently around the head in animals grown at 23^°^C. Occasional animals with such bulges grew slower, presumably due to compromised feeding. In this experiment GFP lethality = 0.8% (4/453), WT lethality = 0.6% (4/671).

**Supplementary Figure 8.**
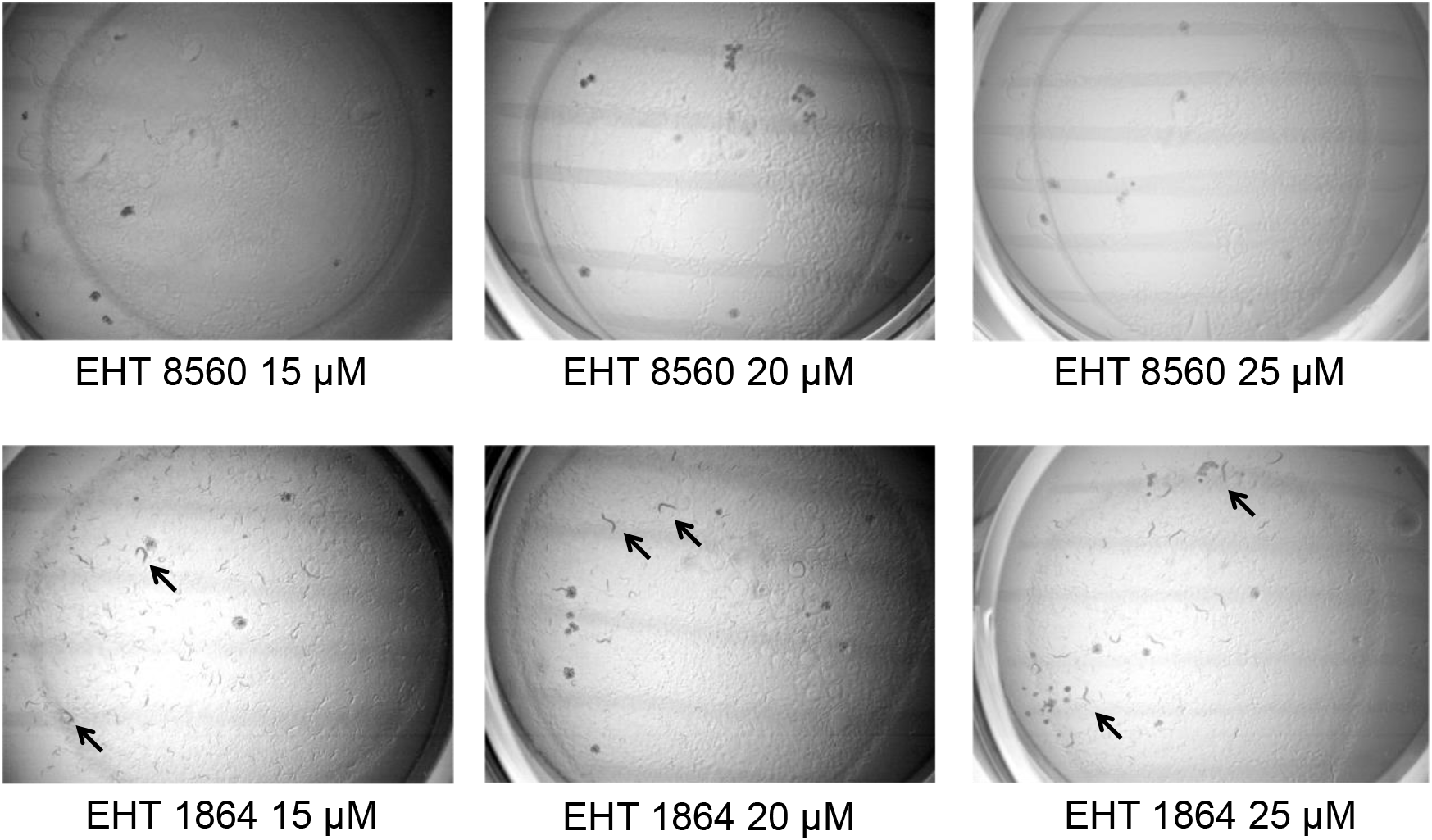
EHT 1864 rescue vs. negative control EHT 8560. 6-well plate assays for small molecule rescue of lethality conferred by *smg-1(cc546*ts); *reIs6* lethality at 23^°^C. **Top row:** Synchronized groups of animals treated with increasing doses of negative control molecule EHT 8560. None survive to hatch. (Tracks are left by Rol parents that laid the eggs). **Bottom row:** Synchronized groups of animals treated with increasing doses of Rac inhibitor EHT 1864. Arrows indicate a subset of grown adults, but many smaller larvae are evident. In all images, dark spots are salt crystals in the agar.

**Supplementary Table 1:** Lethality conferred by component reagents.

| Genotype | 15°C | 23°C |
| --- | --- | --- |
| *smg-1(cc546ts); rels8[P<sub>lin-26</sub>::GFP::NMD<sup>S</sup> 3'UTR]* | 1.9%<br>(4/216) | 1.4%<br>(6/423) |
| *smg-1(cc546ts)* | 0.9%<br>(3/337) | 2.0%<br>(10/522) |
| *rels8[P<sub>lin-26</sub>::GFP::NMD<sup>S</sup> 3'UTR]* | 1.1%<br>(3/284) | 1.2%<br>(5/431) |

**Supplementary Table 2:** EGL-1/BH3-only-induced lethality is caspase-dependent.

| Genotype | RNAi | % lethality<sup>a</sup> |
| --- | --- | --- |
| *smg-1(cc546ts); rels14[P<sub>lin-26</sub>::egl-1(+):NMD<sup>S</sup> 3'UTR]* | *Luc.* | 98.5 (321/326) |
| *smg-1(cc546ts); rels14[P<sub>lin-26</sub>::egl-1(+):NMD<sup>S</sup> 3'UTR]* | *ced-3* | 84.8 (245/289) |
| *smg-1(cc546ts); ced-3(n717); rels14[P<sub>lin-26</sub>::egl-1(+):NMD<sup>S</sup> 3'UTR]* | *Luc.* | 11.4 (22/193) |

<sup>a</sup>Animals were grown at 25°C.

**Supplementary Table 3:** *C. elegans* strains used in this study.

| Strain # | Genotype |
| --- | --- |
| DV2135 | *rels6* [*P<sub>lin-26</sub>::ced-10(gf)::NMD<sup>S</sup> 3'UTR+P<sub>myo-2</sub>::gfp+rol-6(d)*] III |
| DV2149 | *smg-1(cc546ts)* I; *rels6* [*P<sub>lin-26</sub>::ced-10(gf)::NMD<sup>S</sup> 3'UTR+P<sub>myo-2</sub>::gfp+rol-6(d)*] III |
| DV2271 | *smg-1(cc546ts)* I; *max-2(nv162)* II; *rels6* [*P<sub>lin-26</sub>::ced-10(gf)::NMD<sup>S</sup> 3'UTR+P<sub>myo-2</sub>::gfp+rol-6(d)*] III |
| DV2272 | *smg-1(cc546ts)* I; *max-2(nv162)* II; *rels6* [*P<sub>lin-26</sub>::ced-10(gf)::NMD<sup>S</sup> 3'UTR+P<sub>myo-2</sub>::gfp+rol-6(d)*] III |
| DV2286 | *smg-1(cc546ts)* I; *rels6* [*P<sub>lin-26</sub>::ced-10(gf)::NMD<sup>S</sup> 3'UTR+P<sub>myo-2</sub>::gfp+rol-6(d)*] III; *unc115(ky275)* X |
| DV2314 | *smg-1(cc546ts)* *pes-7(gk123)* I; *rels6* [*P<sub>lin-26</sub>::ced-10(gf)::NMD<sup>S</sup> 3'UTR+P<sub>myo-2</sub>::gfp+rol-6(d)*] III |
| DV2316 | *smg-1(cc546ts)* I; *pkn-1(ok1673)* X; *rels6* [*P<sub>lin-26</sub>::ced-10(gf)::NMD<sup>S</sup> 3'UTR+P<sub>myo-2</sub>::gfp+rol-6(d)*] III |
| DV2157 | *smg-1(cc546ts)* *unc-54(r293)* I |
| DV2196 | *smg-1(re1)* *unc-54(r293)* I |
| PD8117 | *smg-1(cc545ts)* *unc-54(r293)* I |
| DV2652 | *smg-1(r861)* *unc-54(r293)* I |
| DV2208 | *unc-97(su110)* X |
| DV2653 | *smg-1(r861)* I; *unc-97(su110)* X |
| DV2654 | *smg-1(cc545ts)* I; *unc-97(su110)* X |
| DV2658 | *smg-1(cc546ts)* I; *unc-97(su110)* X |
| HE110 | *smg-1(re1)* I; *unc-97(su110)* X |
| DV2208 | *unc-97(su110)* X 4x outcrossed |
| DV2471 | *smg-1(cc545ts)* I 2x outcrossed to DV2453 |
| DV2376 | *rels14* [*P<sub>lin-26</sub>::egl-1::NMD<sup>S</sup> 3'UTR + rol-6(d) + P<sub>myo-2</sub>::gfp*] 4x outcrossed |
| DV2437 | *smg-1(cc546ts)* I; *rels14* [*P<sub>lin-26</sub>::egl-1::NMD<sup>S</sup> 3'UTR + rol-6(d) + P<sub>myo-2</sub>::gfp*] |
| DV2348 | *smg-1(cc546ts)* I; *rels8* [*P<sub>lin-26</sub>::gfp::NMD<sup>S</sup> 3'UTR + rol-6(d)*] 4x backcrossed to *smg-1(cc546ts)* |
| DV2453 | *unc-87(e1459)* I 2x outcrossed |
| PD8120 | *smg-1(cc546ts)* I |
| PD8119 | *smg-1(cc545ts)* I |
| DV2683 | *smg-1(cc546ts)* I; *ced-3(n717)* IV; *rels14* [*P<sub>lin-26</sub>::egl-1(+):NMD<sup>S</sup> 3'UTR*] |

**Supplementary Table 4:** Plasmids used in this study.

| Plasmid name | Description |
| --- | --- |
| pML433 | *lin-26(eFGHi) enhancers::minimal myo-2 promoter::GFP* |
| pPD118.44 | *Synthetic NMD<sup>S</sup> 3'UTR* (from inverted let-858 coding sequence) |
| pCM1.3 | *P<sub>lin-26</sub>::synthetic let-858 NMD<sup>S</sup> 3'UTR* |
| pCM1.4 | *P<sub>lin-26</sub>::gfp::synthetic let-858 NMD<sup>S</sup> 3'UTR* |
| pCM3.2 | *P<sub>lin-26</sub>::ced-10(+)::synthetic let-858 NMD<sup>S</sup> 3'UTR* |
| pCM3.3 | *P<sub>lin-26</sub>::ced-10(gf)::synthetic let-858 NMD<sup>S</sup> 3'UTR* |
| pCM12.2 | *P<sub>lin-26</sub>::egl-1::synthetic let-858 NMD<sup>S</sup> 3'UTR* |

**Supplementary Table 5:**
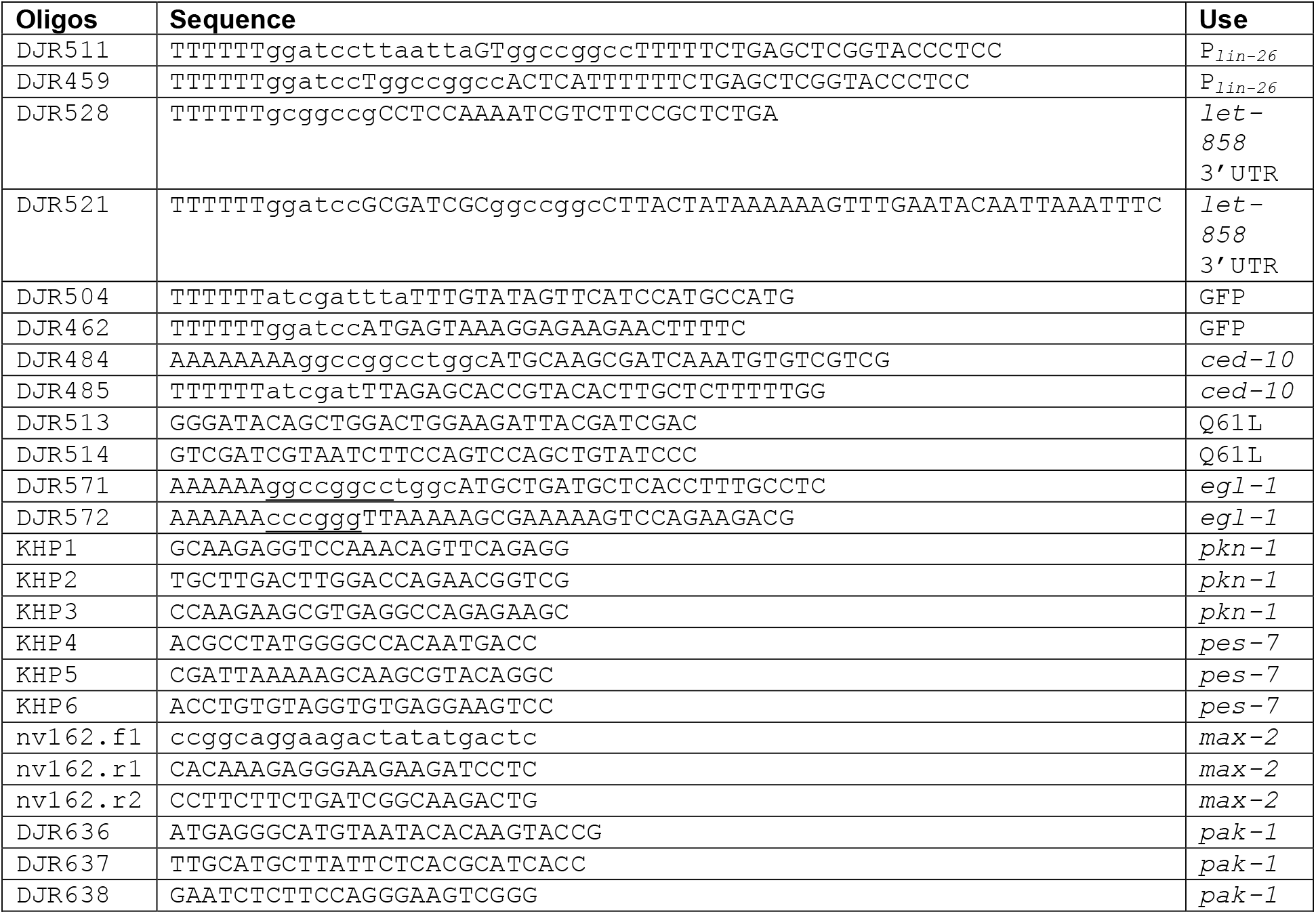
Oligonucleotides used in this study.

## Notes

### Competing Interest Statement

The authors have declared no competing interest.

